# Quantifying Aphantasia through drawing: Those without visual imagery show deficits in object but not spatial memory

**DOI:** 10.1101/865576

**Authors:** Wilma A. Bainbridge, Zoë Pounder, Alison F. Eardley, Chris I. Baker

## Abstract

Congenital aphantasia is a recently characterized experience defined by the inability to form voluntary visual imagery, in spite of intact semantic memory, recognition memory, and visual perception. Because of this specific deficit to visual imagery, aphantasia serves as an ideal population for probing the nature of representations in visual memory, particularly the interplay of object, spatial, and symbolic information. Here, we conducted a large-scale online study of aphantasics and revealed a dissociation in object and spatial content in their memory representations. Sixty-one aphantasics and matched controls with typical imagery studied real-world scene images, and were asked to draw them from memory, and then later copy them during a matched perceptual condition. Drawings were objectively quantified by 2,795 online scorers for object and spatial details. Aphantasics recalled significantly fewer objects than controls, with less color in their drawings, and an increased reliance on verbal scaffolding. However, aphantasics showed incredibly high spatial accuracy, equivalent to controls, and made significantly fewer memory errors. These differences between groups only manifested during recall, with no differences between groups during the matched perceptual condition. This object-specific memory impairment in aphantasics provides evidence for separate systems in memory that support object versus spatial information.

## Introduction

Visual imagery, the ability to form visual mental representations, is a common human cognitive experience, yet it is has been hard to characterize and quantify. What is the nature of the images that come to mind when forming visual representations of objects or scenes? What might these representations look like if one lacks this ability? *Aphantasia* is a recently characterized experience, defined by an inability to create voluntary visual mental images, although semantic memory and vision remain intact (Zeman, Dewar, & Della Sala, 2015; Keogh & Pearson, 2017). Aphantasia is still largely uncharacterized, with many of its studies based on case studies or employing small sample sizes. Here, using an online crowd-sourced drawing task designed to quantify the content of visual memories (Bainbridge, Hall, & Baker, 2019), we examine the nature of aphantasics’ mental representations of visual images within a large sample, and reveal evidence for separate object and spatial systems in human imagery.

Although some cases reporting an absence of mental imagery were first identified in the 19^th^ century (Galton, 1880), the term *aphantasia* has only recently been defined and investigated, within fewer than a dozen studies (Zeman et al., 2015; Keogh & Pearson, 2017; Jacobs, Schwarzkopf, & Silvanto, 2018; Brons, 2019; Wicken, Keogh, Pearson, Unpublished results). This is arguably because most individuals with aphantasia can lead functional, professional lives, with many individuals realizing their imagery experience differed from the majority only in adulthood. The current method for identifying if an individual has aphantasia is through subjective self-report, using the Vividness of Visual Imagery Questionnaire (Marks, 1973). However, recent research has begun quantifying the experience using objective measures such as priming during binocular rivalry (Keogh & Pearson, 2017) and skin conductance during reading (Wicken et al., Unpublished results). Since its identification, several prominent figures have come forth describing their experience with aphantasia, including economics professor Nicholas Watkins (Watkins, 2018), Firefox co-creator Blake Ross (Ross, 2016), and former Pixar Chief Technology Officer Ed Catmull (Gallagher, 2019), leading to broader recognition of the experience.

Like congenital prosopagnosia (Behrmann & Avidan, 2005), in the absence of any brain damage or trauma, aphantasia is considered to be congenital (although it can also be acquired through trauma; Zeman et al., 2010; Thorudottir et al., 2020). However, beyond this, little research has examined the nature of aphantasia and the impact on imagery function and cognition more broadly. A single-participant aphantasia case study found no significant difference from controls in a visual imagery task (judging the location of a target in relation to an imagined shape) nor its matched version of a working memory task, except at the hardest level of difficulty (Jacobs et al., 2018). However, aphantasics show significantly less imagery-based priming in a binocular rivalry task (Keogh & Pearson, 2017; Pearson, 2019), and show diminished physiological responses to fearful text as compared with controls (Wicken et al., Unpublished results). While these studies have observed differences between aphantasics and controls, the nature of aphantasics’ mental representations during visual recall is still unknown. Understanding these differences in representation between aphantasics and controls could shed light on broader questions of what information (visual, semantic, spatial) makes up a memory, and how this information compares to the initial perceptual trace. In fact, the existence of aphantasia serves as key evidence against the hypothesis that visual perception and imagery rely upon the same neural substrates and representations (Dijkstra, Bosch, & van Gerven, 2019), and also suggests a dissociation of visual recognition and recall (as aphantasia only affects the latter). Examination into aphantasia thus has wide-reaching potential implications for the understanding of the way we form mental representations of our world.

The nature and content of our visual imagery has proved incredibly difficult to quantify. Several studies in psychology have developed tasks to objectively study the cognitive process of mental imagery through visual working memory or priming (e.g., Marks, 1973; Marmor & Zaback, 1976; Brandt & Stark, 1997). One of the long-standing debates within the imagery literature has been over the nature of images, and specifically whether visual imagery representations are depictive and picture-like in nature (Kosslyn, 1980; Kosslyn 2005) or symbolic, “propositional” representations (Pylyshyn, 1981; Pylyshyn, 2003). Neuropsychological research, especially in neuroimaging, has led to large leaps in our understanding of visual imagery. Studies examining the role and activation of the primary visual cortex during imagery tasks have been interpreted as supporting the depictive nature of imagery (Ishai, Ungerleider, & Haxby, 2000; Kosslyn, Ganis, & Thompson, 2001; Schacter et al., 2012; Pearson & Kosslyn, 2015). However, neuropsychological studies have identified patients with dissociable impairments in perception versus imagery (Behrmann, 2000; Bartolomeo, 2008), and recent neuroimaging work has suggested there may be systematically related yet separate cortical areas for perception and imagery, and that the neural representation during recall may lack much of the richer, elaborative processing of the initial perceptual trace (Xiao et al., 2017; Silson et al., 2019; Bainbridge, Hall, & Baker, Unpublished results). Combined with research identifying situations where propositional encoding dominates spatial imagery (e.g., Stevens & Coupe, 1978), researchers have concluded that there is a role for both propositional and depictive elements in the imagery process (e.g., Denis & Cocude, 1989). In their case study, Jacobs and colleagues (2018) argue that differences in performance between aphantasic participant AI and neurotypical controls may result from different strategies, including a heavier reliance on propositional encoding, relying on a spatial or verbal code. Thus, ideally a task that measures both depictive (visual) and propositional (semantic) elements of a mental representation could directly compare the strategies used by aphantasics and controls. In a recent study, impressive levels of both object and spatial detail could be quantified by drawings made by neurotypical adults in a drawing-based visual memory experiment (Bainbridge et al., 2019). Such drawings allow a more direct look at the information within one’s mental representation of a visual image, in contrast to verbal descriptions or recognition-based tasks. Thus, a drawing task may allow us to identify what fundamental differences exist between aphantasics and individuals with typical imagery, and in turn inform us of what information exists within imagery.

In the current study, we examine the visual memory representations of congenital aphantasics and individuals with typical imagery (controls) for real-world scene images. Through online crowd-sourcing, we leverage the power of the internet to identify and recruit large numbers of both aphantasic and controls for a memory drawing task. We also recruit over 2,700 online scorers to objectively quantify these drawings for object details, spatial details, and errors in the drawings. We discover a selective impairment in aphantasics for object memory, with significantly fewer visual details and evidence for increased semantic scaffolding. In contrast, for the items that they remember, aphantasics show spatial accuracy at the same high level of precision as controls. Aphantasics also show fewer memory errors and memory correction as compared to controls. These results may point to two systems that support object information versus spatial information in memory.

## Materials and Methods

### Participants

N=123 adults participated in the main online drawing recall experiment, while 2,795 adults participated in online scoring experiments on Amazon Mechanical Turk (AMT) of the drawings from the main experiment. Aphantasic participants for the main experiment were recruited from aphantasia-specific online forums, including “Aphantasia (Non-Imager/Mental Blindness) Awareness Group”, “Aphantasia!” and Aphantasia discussion pages on Reddit. Control participants for the main experiment were recruited from the population at the University of Westminster, online social media sites such as Facebook and Twitter pages for the University of Westminster Psychology, and “Participate in research” pages on Reddit. Scoring participants were recruited from the general population of AMT.

Aphantasic participants were identified by their score on the Vividness of Visual Imagery Questionnaire (VVIQ), a self-report measure of the vividness of one’s visual mental images (Marks, 1973). The VVIQ currently serves as the main tool for identifying aphantasia (although there are other methods, see Keogh & Pearson, 2017). Scores on the VVIQ range from 16 to 80, with aphantasics defined by VVIQ scores ≤ 25 and control VVIQ scores ≥ 40. Eight participants were removed from the analyses for having scores between 26 and 39, and two participants were removed for skipping more than ¼ of the questions in the VVIQ. Of the remaining participants, there were 61 aphantasics and 52 control participants.

No personally identifiable information was collected from any participants, and participants had to acknowledge participation in order to continue, following the guidelines approved by the University of Westminster Psychology Ethics Committee (ETH1718-2345) and the National Institutes of Health Office of Human Subjects Research Protections (18-NIMH-00696).

### Main Experiment: Drawing Recall Experiment

The Drawing Recall Experiment was a fully online memory experiment that consisted of five sections ordered: 1) study phase, 2) recall drawing phase, 3) recognition phase, 4) copied drawing (perception) phase, and 5) questionnaires and demographics. The methods of the experiment are summarized in Fig. 1.

**Fig. 1.**
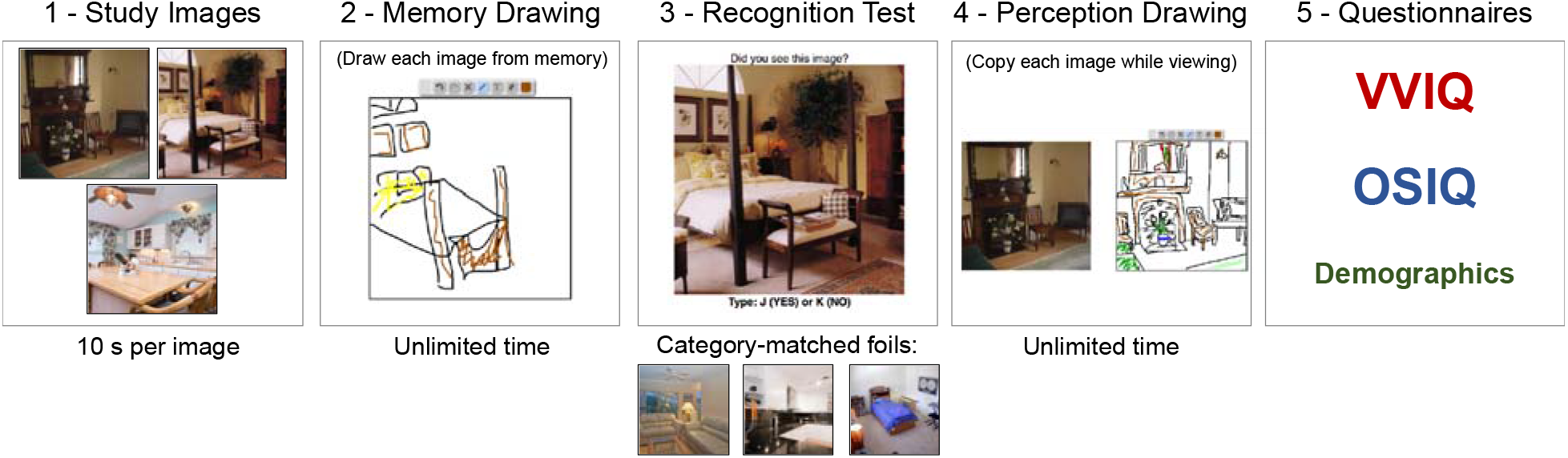
The experimental design of the online experiment. Participants 1) studied three separate scene photographs presented sequentially, 2) drew them from memory, 3) completed a recognition task, 4) copied the images while viewing them, and then 5) filled out the VVIQ and OSIQ questionnaires in addition to demographics questions. The whole experiment took approximately 30 minutes.

First, for the study phase, participants were told to study three images in as much detail as possible. The images were presented at 500 x 500 pixels. They were shown each image for 10 s, presented in a randomized order with a 1 s interstimulus interval (ISI). These three images (see Fig 1a) were selected from a previously validated memory drawing study (Bainbridge et al., 2019), as the images with the highest recall success, highest number of objects, and several unique elements compared to a canonical representation of its category. For example, the kitchen scene does not include several typical kitchen components such as a refrigerator, microwave, or stove, and does include more idiosyncratic objects such as a ceramic chef, zebra-printed chairs, and a ceiling fan. This is important as we want to assess the ability to recall unique visual information beyond just a coding of the category name (e.g., just drawing a typical kitchen). Participants were not informed what they would do after studying the images, to prevent targeted memory strategies.

Second, the recall drawing phase tested what visual memory representations participants had for these images through drawing. Participants were presented with a blank square with the same dimensions as the original images and told to draw an image from memory in as much detail as possible using their mouse. Participants drew using an interface like a simple paint program. They could draw with a pen in multiple colors, erase lines, and undo or redo actions. They were given unlimited time and could draw the images in any order. They were also instructed that they could label any unclear items. Once a participant finished a drawing, they then moved onto another blank square to start a new drawing. They were asked to create three drawings from memory, and could not go back to edit previous drawings. As they were drawing, their mouse movements were recorded to track timing and erasing behavior. These drawings were later quantified by online scorers in a series of separate experiments (see Online Scoring Experiments below).

Third, the recognition phase tested whether there was visual recognition memory for these specific images. Participants viewed images and were told to indicate whether they had seen each image before or not. The images consisted of the three images presented in the study phase as well as three new foil images of the same scene categories (kitchen, bedroom, living room). Matched foils were used so that recognition performance could not rely on recognizing the category type alone. All images were presented at 500 x 500 pixels. Participants were given unlimited time to view the image and respond, and a fixation cross appeared between each image for 200 ms.

Fourth, the copied drawing phase had participants copy the drawings while viewing them, in order to see how participants perceive each image in the absence of a memory task. This phase gives us an estimate of the participant’s drawing ability and ability to use this drawing interface with a computer mouse to create drawings. This phase also measures the maximum information one might draw for a given image (e.g., you won’t draw every plate stacked in a cupboard). Participants saw each image from the study phase presented next to a blank square. They were instructed to copy the image in as much detail as possible. The blank square used the same interface as the recall drawing phase. When they were done, they could continue onto the next image, until they copied all three images from the study phase. The images were tested in a random order, and participants had as much time as they wanted to draw each image, but could not go back to any completed drawings.

Finally, participants filled out three questionnaires at the end. They completed the previously mentioned VVIQ (Marks, 1973), which was mainly used to determine participant group membership. Participants also completed the more recent Object and Spatial Imagery Questionnaire (OSIQ) (Blajenkova, Kozhevnikov, & Motes, 2006), which measures visual imagery preference for object information and spatial information, providing a score between 15-75 for each subscore (object, spatial). Finally, participants provided basic demographics, basic information about their computer interface, and their experience with art. In these final questions, they indicated which component of the experiment was most difficult, and were able to write comments on why they found it difficult.

### Online Scoring Experiments

In order to objectively and rapidly score the 692 drawings produced in the Drawing Recall Experiment, we conducted online crowd-sourced scoring experiments with a set of 2,795 participants on AMT, an online platform used for crowd-sourcing of tasks. None of these participants took part in the Drawing Recall Experiment. For all online scoring experiments, scorers could participate in as many trials as they wanted, and were compensated for their time.

#### Object Selection Study

AMT scorers were asked to indicate which objects from the original images were in each drawing. This allows us to systematically measure how many and what types of objects exist in the drawings. They were presented with one drawing and five photographs of the original image with a different object highlighted in red. They had to click on all object images that were contained in the original drawing. Five scorers were recruited per object, with 909 unique scorers in total. An object was determined to exist in the drawing if at least 3 out of 5 scorers selected it.

#### Object Location Study

For each object, AMT scorers were asked to place and resize an oval around that object in the drawing, in order to get information on the location and size accuracy of the objects in the drawings. AMT scorers were instructed on which object to circle in the drawing by the original image with the object highlighted in red, and only objects selected in the Object Selection Study were used. Five scorers were recruited per object, with 1,310 unique scorers in total. Object location and size (in both the x and y directions) were taken as the median pixel values across the five scorers.

#### Object Details Study

AMT scorers here indicated what details existed in the specific drawings. In a first AMT experiment, five scorers per object (N=304 total) saw each object from the original images and were asked to list 5 unique traits about the object (e.g., shape, material, pattern, style). A list of unique traits was then created for each object in the images. In a second AMT experiment, scorers were then shown each object in the drawings (highlighted by the ellipse drawn in the Object Location Study), and had to indicate whether that trait described the drawn object or not. Five scorers were recruited per trait per drawn object, with 777 unique scorers in total.

#### False Objects Study

AMT scorers were asked to indicate “false objects” in the drawings—what objects were drawn in the drawing that didn’t exist in the original image? Scorers were shown a drawing and its corresponding image and were asked to write down a list of all false objects. Nine scorers were recruited per drawing, with 337 unique scorers in total. An object was counted as a false object if at least three scorers listed it.

### Additional Drawing Scoring Metrics and Analyses

In addition to the Online Scoring Experiments, other attributes were collected for the drawings. A blind scorer (the corresponding author) went through each drawing presented in a random order (without participant or condition information visible) and had to code *yes* or *no* for if the drawing 1) contained any color, 2) contained any text, and 3) contained any erasures. Erasures were quantified by viewing the mouse movements used for drawing the image, to see if lines were drawn and then erased, and did not make it into the final image.

Throughout this manuscript, whenever parametric statistical tests were used to compare groups, we first confirmed the measures were not significantly different from a normal distribution, using the Kolmogorov-Smirnov test of goodness-of-fit.

## Results

With these memory and perceptual drawings, we can then make direct comparisons in the types of detail, amounts of detail, and types of errors that may differ between aphantasics and controls. First, we examine the demographic measures between the two groups, such as age, gender, art ability, and ratings on the OSIQ. Second, we turn to objective quantification of the drawings, and explore differences in the objects drawn by aphantasics and controls and text-based strategies. Third, we compare spatial accuracy in the drawings between these two groups. Finally, we compare the presence of memory errors, quantifying the number of falsely inserted additional objects.

### No demographic differences between groups, but reported differences in object and spatial imagery

First, we analyzed whether there were demographic differences between the groups. There was a significant difference in age between groups with aphantasics generally older than controls (aphantasic: *M*=41.88 years, *SD*=13.88, *Range*=18 to 74 years; control: *M*=32.12 years, *SD*=15.26, *Range*=18 to 75 years; *t*(107)=3.49, *p*=6.95 × 10^−4^). There was no significant difference in gender proportion between the two groups (aphantasic: 62.3% female; control: 59.6% female; Pearson’s chi-square test for proportions: *χ*^2^ =0.08, p=0.771), even though a previous study reported a sample comprising of predominantly males (Zeman et al., 2015).

Second, we investigated the relationship of the VVIQ score and OSIQ (Fig. 2), a questionnaire developed to separate abilities to perform imagery with individual objects versus spatial relations amongst objects (Blajenkova et al., 2006). Controls scored significantly higher on the OSIQ than aphantasics (*t*(103) = 12.70, *p*=8.55 × 10^−23^). There was a significant correlation between VVIQ score and OSIQ score for control participants (*M*=89.73, *SD*=10.97; Spearman rank-correlation test: *ρ*=0.54, *p*=7.70 × 10^−5^), but only marginally for aphantasics (OSIQ M score=62.88, SD=10.65; ρ=0.26, p=0.052). When broken down by OSIQ subscale, there was a significant difference between groups in questions relating to object imagery (*t*(103)=20.00, p=3.01 × 10), but not spatial imagery (*t*(103)=-0.33, *p*=0.742). Indeed, a 2-way ANOVA (participant group × subscale) reveals a main effect of participant group (*F*(1,206)=154.97, *p*~0), subscale (*F*(1,206)=40.11, *p*=1.48 × 10^−9^), and a significant interaction (*F*(1,206)=167.94, *p*~0), confirming a difference in self-reported ratings for object imagery and spatial imagery respectively. This difference in self-reported object imagery and spatial imagery has been reported a previous study (Keogh & Pearson, 2017), and suggests a potential difference between the two imagery subsystems

**Figure 2.**
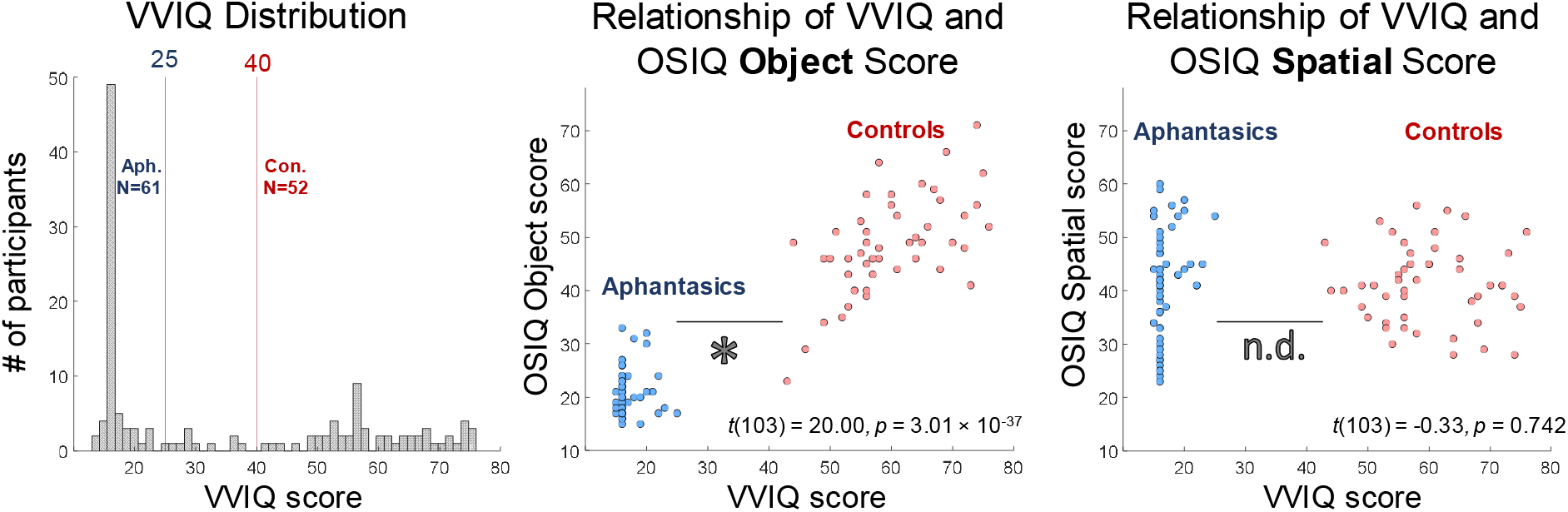
Experimental paradigm and basic demographics. a) b) (Left) A histogram of the distribution of participants across the VVIQ. Aphantasics were selected as those scoring 25 and below (N=61) and controls were selected as those scoring 40 and above (N=52), while those in between were removed from the analyses (N=8). While the range of the VVIQ is from 16 to 80, some participants (N=10 out of 121 total) skipped 1-3 questions, leading to some participants scoring below 16. These skipped questions did not affect group membership. (Middle) A scatterplot of total VVIQ score plotted against total OSIQ Object component score for participants meeting criterion. Each point represents a participant, with aphantasics in blue and controls in red. There was a significant difference in OSIQ Object score between the two groups. (Right) A scatterplot of total VVIQ score plotted against OSIQ Spatial component score. There was no difference in OSIQ Spatial score between the two groups. Both the OSIQ Object component and Spatial components have a range of 15 to 75 points.

Third, we investigated whether aphantasics and controls reported different levels of comfort or familiarity with art, which may influence their drawing performance. When asked to rate their artistic abilities on a scale from 1 (very poor) to 5 (very good), aphantasics and controls showed no significant difference in their ratings (aphantasic: *M*=2.30, *SD*=1.34; control: *M*=2.52, *SD*=0.99; non-parametric Wilcoxon rank sum test: *Z*=1.23, *p*=0.219). Both aphantasic and control participants also reported taking art classes in the past (39.34% of aphantasic participants, 37.74% of controls). Many aphantasics (13.11%) also reported being employed within industries involving artistic abilities, such as sculpting, visual arts, makeup art, and interior decoration. Thus, aphantasics and controls did not show strong differences in their propensity for, or interest in, art.

Finally, given the focus of the current experiment on visual recall, we also compared measures of visual recognition performance. Both groups performed near ceiling at visual recognition of the images they studied, with no significant difference between groups in recognition hit rate (controls: M=0.96, SD=0.12; aphantasics: M=0.97, SD=0.12; Wilcoxon rank-sum test: *Z*=1.09, *p*=0.274), or false alarm rate (controls: M=0.02, SD=0.12; aphantasics: M=0, SD=0; Wilcoxon rank-sum test: *Z*=1.10, *p*=0.273). These results indicate that there is no obvious deficit in aphantasics for recognizing images, even with lures from the same semantic scene category.

### Diminished object information for aphantasics

Next, we turned to analyzing the drawings made by the participants to reveal objective measures of the mental representations of these two groups. Looking at overall number of drawings made, while a small number of participants could not recall all three images, there was no significant difference between groups in number of images drawn from memory (control: *M*=2.92, *SD*=0.27; aphantasic: *M*=2.89, *SD*=0.37; Wilcoxon rank-sum test: *Z*=0.42, *p*=0.678). To evaluate the drawings, 2,795 unique workers from the online experimental platform Amazon Mechanical Turk (AMT) scored the drawings on a variety of metrics including object information, spatial accuracy, and memory errors, using methods previously established for quantifying memory drawings (Bainbridge et al., 2019). Importantly, each participant completed both a memory drawing (i.e., drawing an image from memory for an unlimited time period) and a perception drawing (i.e., copying from a drawing for an unlimited time period) for each image, allowing us to compare for each participant what is in memory versus what that individual would maximally draw given an image without memory constraints (refer to Fig. 3 for example drawings). This comparison allows us to control for differences in effort and drawing ability, which we should expect to be reflected in both types of drawings.

**Figure 3.**
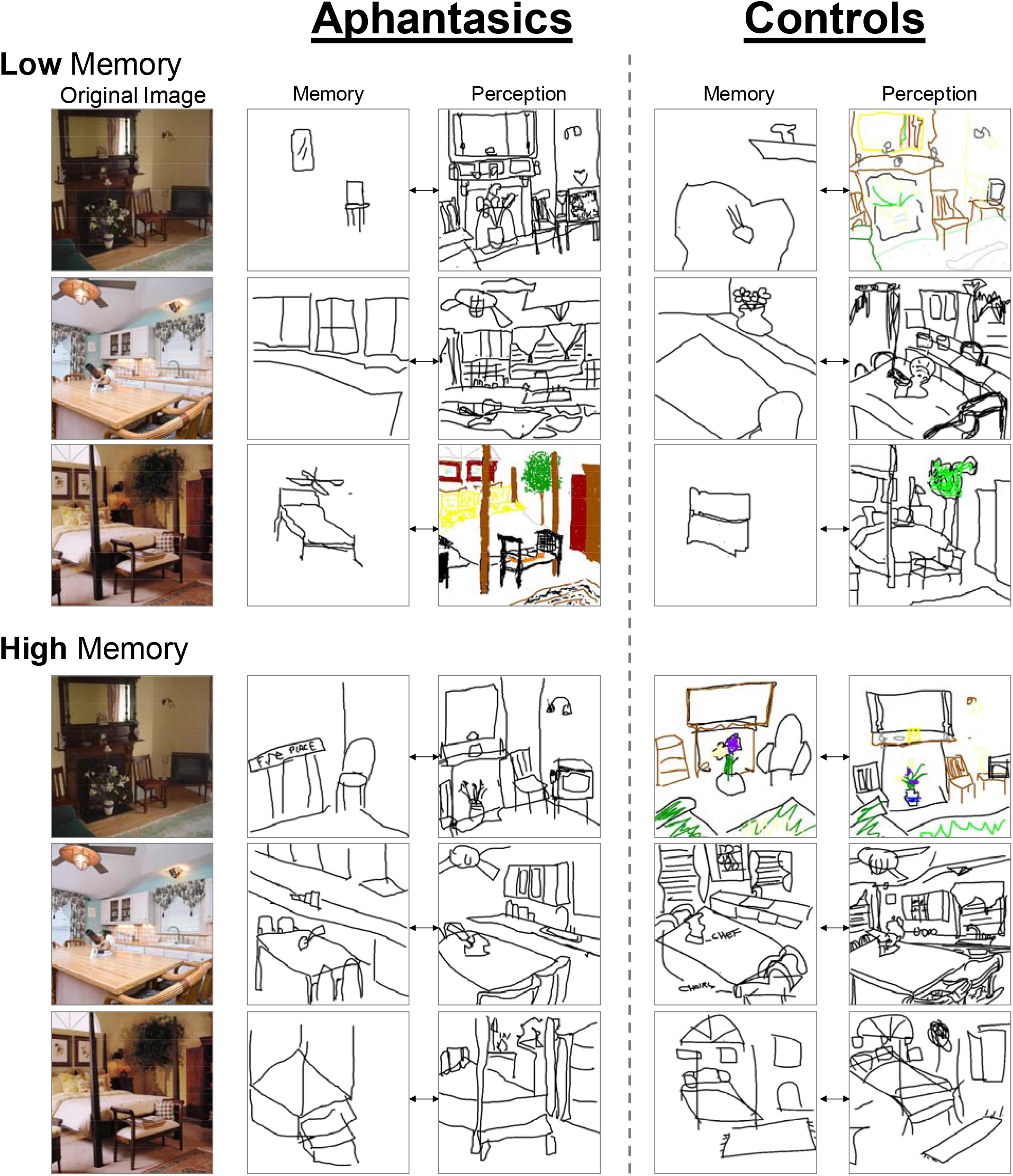
Example drawings. Example drawings made by aphantasic and control participants from memory and perception (i.e., copying the image) showing the range of performance. The memory and perception drawings connected by arrows are from the same participant, and every row is from a different participant. Low memory examples show participants who drew the fewest from memory but the most from perception. High memory examples show participants who drew the highest amounts of detail from both memory and perception. These examples are all highlighted in the scatterplot of Fig. 4. The key question is whether there are meaningful differences between these two sets of participants’ drawings.

To score level of object information, AMT workers (N=5 per object) identified whether each of the objects in an image was present in each drawing of that image (Fig. 4). A 2-way ANOVA of participant group (aphantasic / control) × drawing type (memory / perception drawing, repeated measure) looking at number of objects drawn per image showed no significant overall effect of participant group (*F*(1,223)=0.26, *p*=0.613), but a significant effect of drawing type (*F*(1,223)=507.03, *p*~0), and more importantly, a significant statistical interaction (*F*(1,223)=9.25, *p*=0.0029). Targeted post-hoc independent t-tests revealed that when drawing from memory, controls drew significantly more objects (*M*=6.32 objects per image, *SD*=3.07) than aphantasics (*M*=4.98, *SD*=2.54; *t*(111)=2.53, *p*=0.013) across the experiment. In contrast, when copying a drawing (perception drawing), aphantasics on average drew more objects from the images than controls, but with no significant difference (controls: *M*=18.00 objects per image, *SD*=5.81; aphantasics: *M*=20.07, *SD*=7.26; *t*(111)=1.74, *p*=0.085). These results suggest that aphantasics are showing a specific deficit in recalling object information during memory.

**Figure 4.**
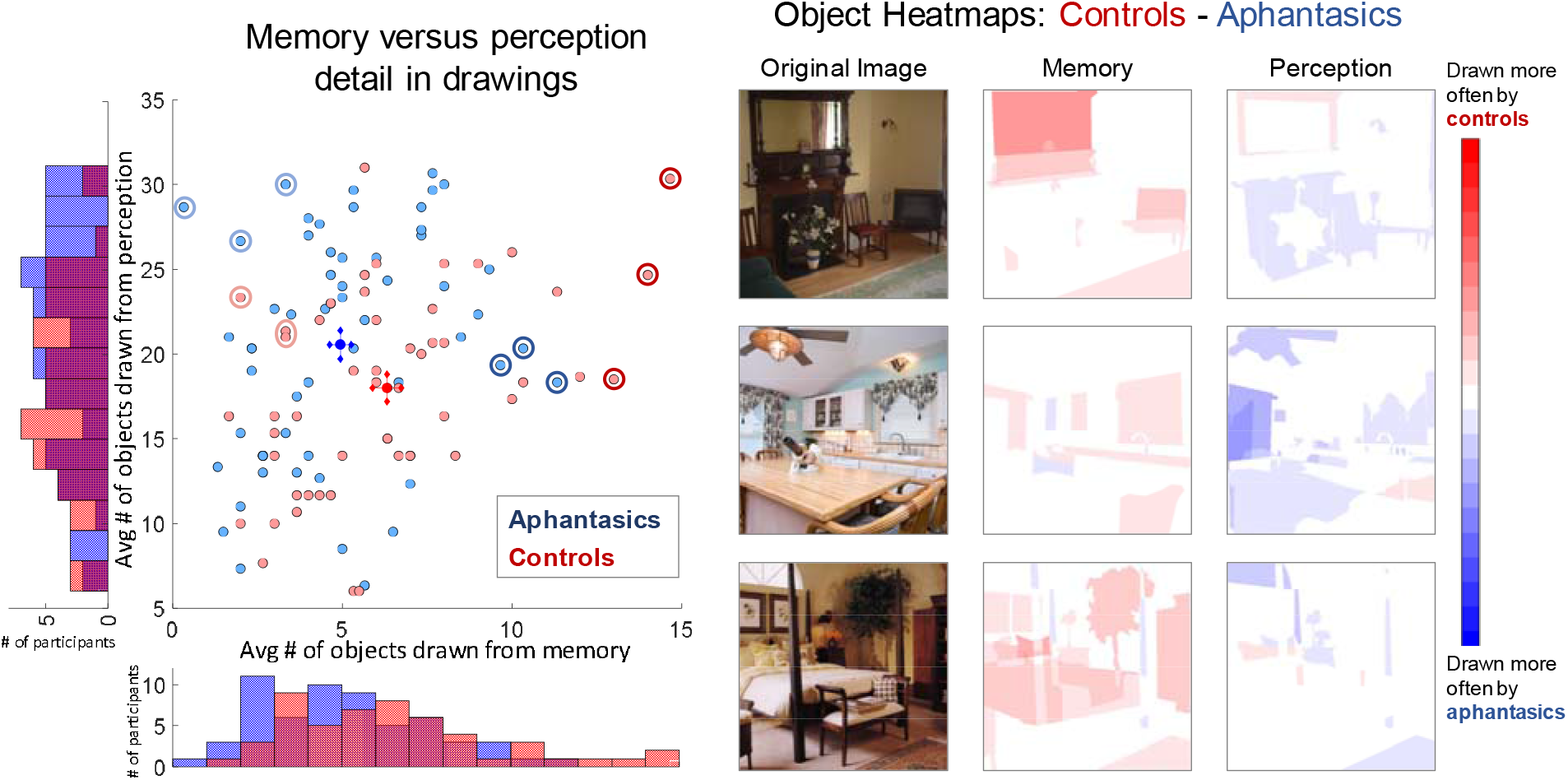
Comparison of object information in drawings between aphantasics and controls. (Left) A scatterplot of each participant as a point, showing average number of objects drawn from memory across the three images (x-axis), versus average number of objects drawn from perception across the three images (y-axis). Aphantasics are in blue, while controls are in red. The bright blue circle indicates average aphantasic performance, while the bright red circle indicates average control performance, with crosshairs for both indicating standard error of the mean for memory and perception respectively. Histograms on the axes show the number of participants who drew each number of objects. Controls drew significantly more objects from memory, although with a tendency towards fewer from perception. The highlighted light blue and red points are the participants with the lowest memory performance shown in Fig. 3, while the highlighted dark blue and red points are the participants with the highest memory performance shown in Fig. 3. (Right) Heatmaps of which objects for each image tended to be drawn more by controls (red) or aphantasics (blue). Pixel value represents the proportion of control participants who drew that object in the image subtracted by the proportion of aphantasics who drew that object (with a range of −1 to 1). Controls remembered more objects (i.e., there is more red in the memory heatmaps), even though aphantasics tended to copy more objects (i.e., there is more blue in the perception heatmaps).

Given that some participants tended to draw few objects even when copying from an image, we also investigated a corrected measure, taken as the number of objects drawn from memory divided by the number of objects drawn from perception, for each image for each participant. Drawings from perception with fewer than 5 objects were not included in the analysis, to remove any low-effort trials. Aphantasics drew a significantly smaller proportion of objects from memory than control participants (aphantasic: *M*=0.261, *SD*=0.165; control: *M*=0.369, *SD*=0.162; Wilcoxon rank-sum test: *Z*=4.09, *p*=4.24 × 10^−5^). We also investigated the correlation within groups between the number of objects drawn from memory and the number drawn from perception. There was a significant correlation for both groups, where the more one draws from perception, the more one also tends to draw from memory (Pearson correlation; aphantasics: *r*=0.34, *p*=0.0075; controls: *r*=0.40, *p*=0.0035).

We also assessed the relationship between performance in the task and self-reported object imagery in the OSIQ. Across groups, there was a significant correlation between proportion of objects drawn from memory and OSIQ object score (Spearman’s rank correlation: *ρ*=0.33, *p*=7.18 × 10^−4^), although these correlations were not significant when separated by participant group (*p*>0.10).

Next, we examined whether there was a difference in visual detail within objects, by quantifying differences between groups in color and amount of time spent on the drawings. Significantly more memory drawings by controls contained color than those by aphantasics (control: 38.2%, aphantasics: 21.6%; Pearson’s chi-square test for proportions: *χ*^2^ =10.09, *p*=0.0015), while there was no significant difference for perception drawings (control: 46.2%, aphantasic: 39.4%, *χ*^2^ =1.46, *p*=0.227). Control participants also spent significantly longer time on their memory drawings than aphantasics (control: *M*= 119.41 s per image, *SD*=68.88 s; aphantasics: *M*=71.22 s, *SD*=49.17 s; *t*(110) = 4.31, *p*=3.56 × 10^−5^), implying more attention to detail in their drawings. For the perception drawings, there was no significant difference between groups in the amount of time they spent on their drawings (control: *M*=272.33 s, *SD*=214.17 s; aphantasic: *M*=295.18 s, *SD*=304.54 s; *p* = 0.654). These differences in time spent on memory drawing could reflect controls spending more time because they drew more objects from memory. However, even if we normalize total drawing time by number of objects drawn to get an estimate of average time spent per object, controls spent significantly more time per object than aphantasics when drawing from memory (Wilcoxon rank sum test: *Z*=2.09, *p*=0.037), but not when drawing during perception (*Z*=0.75, *p*=0.454). This implies that aphantasics not only spent less time per drawing, but also less time on the details for each object. Finally, we investigated other forms of object detail, by having AMT workers (N=777) judge whether different object descriptors (e.g., material, texture, shape, aesthetics; generated by 304 separate AMT workers) applied to each drawn object. This task did not identify differences between groups for the memory drawings (t(110)=0.21, p=0.833), although objects were significantly more detailed when copied than when drawn from memory for both aphantasics (memory: *M*=42.4% descriptors per object applied, SD=5.1%; copied: *M*=45.9%, *SD*=4.1%; *t*(119)=4.12, p=6.92 × 10) and control participants (memory: *M*=42.2%, SD=5.6%; copied: *M*=47.0%, *SD*=3.9%; *t*(100)=5.06, *p*=1.92 × 10). However, it is possible this task may have asked for too fine-grained information than can be measured from these drawings (e.g., judging the material and texture of a drawn chair).

In sum, these results present concrete evidence that aphantasics recall fewer objects than controls, and these objects contain less visual detail (i.e., color, less time spent for drawing) within their memory representations.

### Aphantasics show greater dependence on symbolic representations

While aphantasics show decreased object information in their memory drawings, they are still able to successfully draw some objects from memory (4.98 objects per image on average). Do these drawings reveal evidence for alternative, non-visual strategies that may have supported this level of performance? To test this question, we quantified the amount of text used to label objects included in the participants’ drawings. Note that while labeling was allowed (the instructions stated: “Please draw or label anything you are able to remember”), it was effortful as it required drawing the letters with the mouse. We found that significantly more memory drawings by aphantasics contained text than those by controls (aphantasic: 29.6%, control: 16.0%; *χ*^2^ =7.57, p=0.0059). Further, there was no significant difference between groups for perception drawings (aphantasic: 2.9%, control: 0.8%; *χ*^2^ =1.77, *p*=0.184). These results imply that aphantasics may have relied upon symbolic representations to support their memory.

One question is whether aphantasics just prefer writing over drawing, and so prioritized time or effort on writing text over drawing objects. To elaborate, it is possible that aphantasics expend their effort on writing text, and then do not want to spend further time on drawing objects even if they might have object information in memory. If this were the case, then drawings that contain text should contain fewer objects. However, we found there was no significant difference in number of objects between aphantasic memory drawings with text and without (independent samples t-test by drawing: *t*(174)=0.07, *p*=0.947). There was also no significant difference for their drawings made during perception (*t*(171)=0.35, *p*=0.726), nor were there differences for controls (memory drawings: *t*(150)=0.004, *p*=0.997; perception drawings: *t*(152)=1.50, *p*=0.135). These results indicate that the usage of text was not a trade-off with object memory; aphantasics preferred to include text in their memory drawings regardless of how many objects they recalled.

Comments by aphantasics at the end of the experiment supported their use of symbolic strategies. When asked what they thought was difficult about the task, one participant noted, “Because I don’t have any images in my head, when I was trying to remember the photos, I have to store the pieces as words. I always have to draw from reference photos.” Another aphantasic stated, “I had to remember a list of objects rather than the picture,” and another said, “When I saw the images, I described them to myself and drew from that description, so I…could only hold 7-9 details in memory.” In contrast, control participants largely commented on their lack of confidence in their drawing abilities: e.g., “I am very uncoordinated so making things look right was frustrating”; “I can see the picture in my mind, but I am terrible at drawing.”

### Aphantasics and controls show equally high spatial accuracy in memory

While aphantasics show an impairment in memory for object information, do they also show an impairment in spatial placement of the objects? To test this question, AMT workers (N=5 per object) drew an ellipse around the drawn version of each object, allowing us to quantify the size and location accuracy of each drawn object (Fig. 5). When drawing from memory, there was no significant difference between groups in object location error in the x-direction (aphantasic: *M* pixel error=63.86, *SD*=31.59; control: *M*=60.63, *SD*=28.45; *t*(111)=0.57, *p*=0.572) nor the y-direction (aphantasic: *M*=65.43, *SD*=29.89; control: *M*=69.10, *SD*=29.72; *t*(111)=0.65, *p*=0.515). However, this lack of difference was not due to difficulty in spatial accuracy; both groups’ drawings were incredibly spatially accurate, with all average errors in location less than 10% of the size of the images themselves. Similarly, there was also no significant difference in drawn object size error in terms of width (aphantasic: *M* pixel error=23.00, SD=10.95; control: *M*=24.89, *SD*=13.58; *t*(111)=0.82, *p*=0.413) and height (aphantasic: *M*=26.75; *SD*=14.15; control: *M*=22.82; *SD*=11.05; *t*(111)=1.62, *p*=0.107), and these sizes were incredibly accurate in both groups (average errors less than 4% of the image size). There was no correlation between a participant’s level of object location or size error and ratings on the OSIQ spatial questions (all *p*>0.30). In all, these results show that both aphantasics and controls have highly accurate memories for spatial location, with no observable differences between groups.

**Figure 5.**
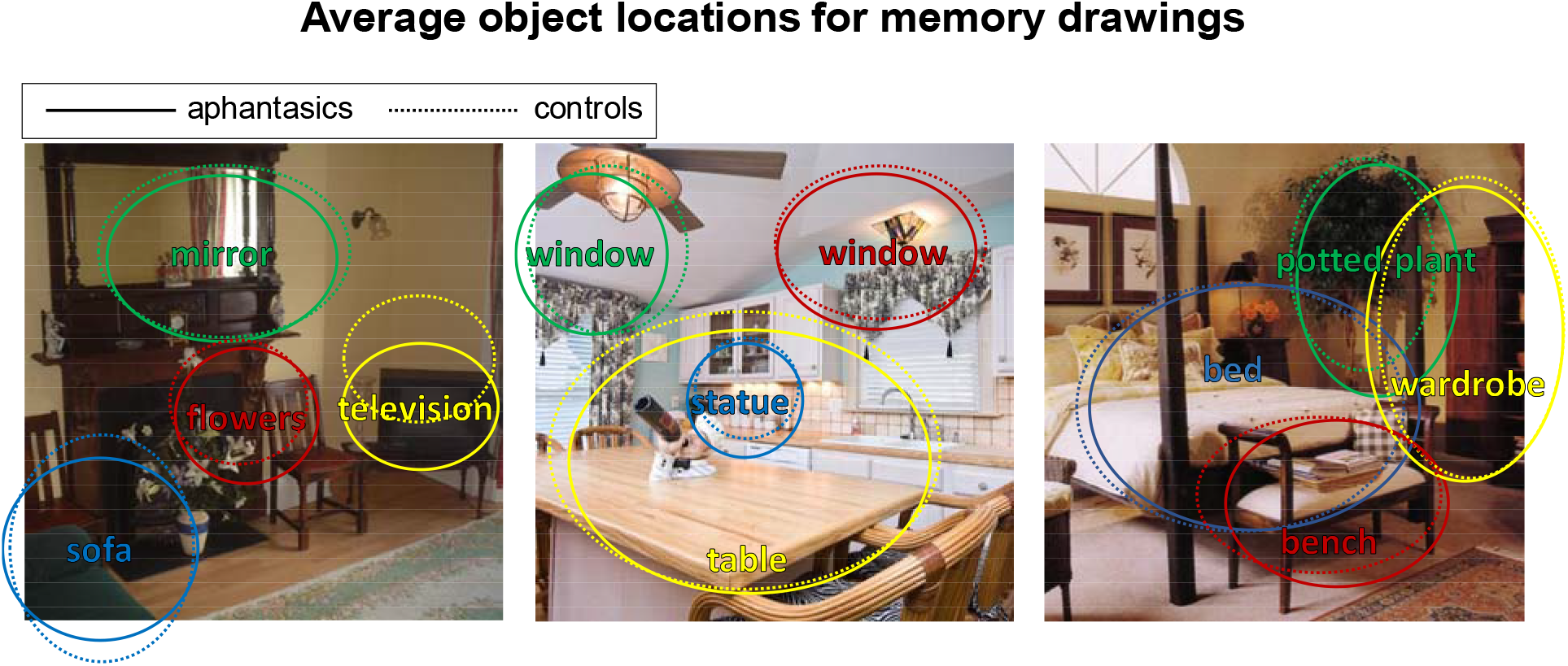
Average object locations and sizes recalled by aphantasics and controls. Average object locations and sizes for memory drawings of four of the main objects from each image, made by aphantasics (solid lines) and controls (dashed lines). Even though these objects were drawn from memory, their location and size accuracy was still very high. Importantly, aphantasics and controls showed no significant differences in object location or size accuracy.

### Aphantasics draw fewer false objects than controls

Finally, we quantified the amount of error in participants’ drawings from memory by group. AMT workers (N=5 per drawing) viewed a drawing and its corresponding image and wrote down all objects in the drawings that were not present in the original image (essentially quantifying false object memories). Significantly more memory drawings by controls contained false objects than drawings by aphantasics (controls: 14 drawings, aphantasics: 3 drawings; Pearson chi-square test: *χ*^2^ =9.35, *p*=0.002); examples can be seen in Fig. 6. This is not just because controls drew more objects overall and were thus more likely to draw false objects. If we also look at proportion of total objects drawn by group that were false objects, significantly more objects drawn by controls were false objects than those drawn by aphantasics (*χ*^2^ =6.37, *p*=0.012). This indicates that control participants were making more memory errors, even after controlling for the fewer number of objects drawn overall by aphantasics. Interestingly, all aphantasic errors (see Fig. 6) were transpositions from another image and drawn in the correct location as the original object (a tree from the bedroom to the living room, a window from the kitchen to the living room, and a ceiling fan from the kitchen to the bedroom). In contrast, several false memories from controls were objects that did not exist across any image but instead appeared to be filled in based on the scene category (e.g., a piano in the living room, a dresser in the bedroom, logs in the living room). No perception drawings by participants from either group contained false objects.

**Figure 6.**
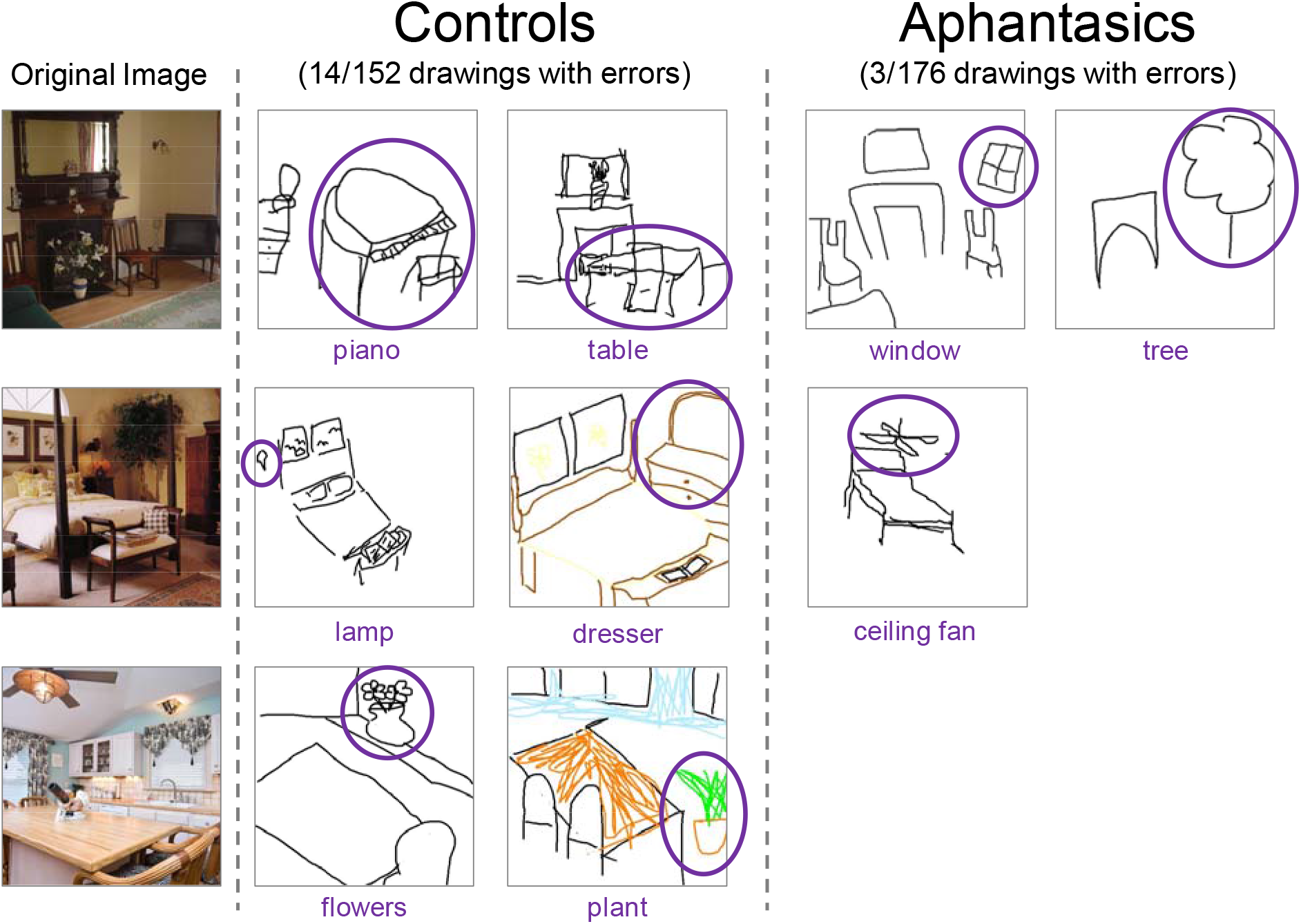
False object memories in the drawings. Examples of the false object memories made by participants in their memory drawings, with the inaccurate objects circled. Control participants made significantly more errors, with only 3 out of 176 total aphantasic drawings containing a falsely remembered object. Note that all aphantasic errors were also transpositions from other images.

As another metric of memory error, we also coded whether a drawing was edited or not, based on tracked mouse movements. A drawing was scored as edited if at least one line was drawn and then erased during the drawing. Significantly more memory drawings by control participants had editing than those by aphantasic participants (aphantasic: 28.4%, control: 46.6%; *χ*^2^ =10.72, *p*=0.0011). There was no significant difference in editing between groups for the perception drawings (aphantasic: 37.6%, control: 47.7%; *χ*^2^ =3.17, *p*=0.075), indicating these differences are likely not due to differences in effort.

## Discussion

Through a drawing task with a large online sample, we conducted an in-depth characterization of the mental representations held by congenital aphantasics, a recently identified group of individuals who self-report the inability to form voluntary visual imagery. We discover that aphantasics show impairments in object memory, drawing fewer objects, containing less color, and spending less time drawing details. Further, we find evidence for greater dependence on symbolic information in the task, with more text in their drawings and common self-reporting of verbal strategies. However, aphantasics show no impairments in spatial memory, positioning objects at accurate locations with the correct sizes. Further, aphantasics show significantly fewer errors in memory, with fewer falsely recalled objects, and less correction of their drawings. Importantly, we observe no significant differences between controls and aphantasics when drawing directly from an image, indicating these differences are specific to memory and not driven by differences in effort, drawing ability, or perceptual processing. Indeed, aphantasics reported an equal confidence in their art abilities compared to controls, and many had experience with art classes and art-based careers.

Collectively, these results point to a dissociation in imagery between object-based information and spatial information. In addition to selective deficits in object memory over spatial memory, aphantasics subjectively report a lower preference for object imagery compared to spatial imagery in the OSIQ. This supports subjective self-report in the smaller dataset (N=15) of Keogh & Pearson (2017), which first reported differences in OSIQ measures. Further, in the current study, participants’ reported object imagery abilities correlated with the number of objects they drew from memory. These consistent results both confirm the OSIQ as a meaningful measure, while also demonstrating how such deficits can be captured by a behavioral measure such as drawing. While a similar dissociation between object and spatial memory has been observed in other paradigms and populations (Farah & Hammond, 1988), the current study provides further evidence for this dissociation in a population of individuals in the absence of trauma or changes in brain pathology. Cognitive decline from aging and dementia have shown selective deficits in object identification versus object localization (Reagh et al., 2016), owing to changes in the medial temporal lobe, where the perirhinal cortex is thought to contribute to object detail recollection, while the parahippocampal cortex contributes to scene detail recollection (Staresina, Duncan, & Davachi, 2011). The neocortex is also considered to be organized along separate visual processing pathways, with ventral regions primarily coding information about visual features, and parietal regions coding spatial information (Farah, Hammond, Levine, & Calvanio, 1988; Ungerleider & Haxby, 1994; Corballis, 1997; Carlesimo, Perri, Turriziani, Tomaiuolo, & Caltagirone, 2001; Kravitz, Saleem, Baker, & Mishkin, 2011). These findings also suggest interesting parallels between the imagery experience of individuals with aphantasia and individuals who are congenitally blind, who perform similarly to typically sighted individuals on a variety of spatial imagery tasks (Kerr, 1983; Zimler & Keenan, 1983; Eardley & Pring, 2007; Cattaneo et al., 2008). Neuroimaging of aphantasics will be an important next step, to see whether these impairments are manifested in decreased volume or connectivity of regions specific to the imagery of visual details, such as anterior regions within inferotemporal cortex (Ishai et al., 2000; O’Craven & Kanwisher, 2000; Lee, Kravitz, & Baker, 2012; Bainbridge et al., Unpublished results) or medial parietal regions implicated in memory recall (Buckner, Andrews-Hanna, & Schacter, 2008; Vilberg & Rugg, 2008; Ranganath & Ritchey, 2012; Silson et al., 2019).

Further investigations on aphantasics will also provide critical insight on the nature of imagery, and how it compares to different forms of memory. While aphantasics show an impairment at recall performance, no evidence has shown impairments in visual recognition, and indeed our study also observes near-ceiling recognition performance. These results support other converging evidence pointing towards a neural dissociation in the processes of quick, automatic visual recognition and slower, elaborative visual recall (Jacoby, 1991; Holdstock et al., 2002; Staresina & Davachi, 2006; Barbeau, Pariente, Felician, & Puel, 2011; Bainbridge et al., 2019). That being said, the recognition task in the current experiment had relatively low difficulty, testing foil images of the same semantic category, but without other matched detail (e.g., identities of objects). Future work could study whether aphantasics are impaired at more fine-grained recognition tasks, where object and spatial detail within an image are selectively manipulated. Aphantasics also report fully intact verbal recall abilities, and our results suggest that they may be using semantic strategies, in combination with accurate spatial representations, to compensate for their lack of visual imagery. In fact, in the current study, aphantasics’ drawings from memory contained more text than those of controls, potentially indicating a semantic propositional coding of their memories to perform the task. Imagery of a visual stimulus thus may not necessarily be visual in nature; while forming a visual representation of the scene or object may be one way to undertake the task, there may be other, non-visual strategies to complete the task. Even in neurotypical adults, imagery-based representations in the brain may differ from perceptual representations of the same items (Bainbridge et al., Unpublished results). Further neuroimaging investigations will lead to an understanding of the neural mechanisms underlying these different strategies.

Further, aphantasics’ lower errors in memory (e.g., fewer falsely recalled objects compared to controls) could possibly reflect higher accuracy in semantic memory versus controls, to compensate for visual memory difficulties. Aphantasics may serve as an ideal population to probe the difference between visual and semantic memory and their interaction in both behavior and the brain. Additionally, while aphantasia has thus far only been quantified in the visual domain, preliminary work suggests that the experience may extend to other modalities (Zeman et al., 2015). Using a multimodal approach, researchers may be able to pinpoint neural differences in aphantasics across other sensory modalities, for instance, the auditory domain which has been shown to have several characteristics similar to the visual domain (Halpern, 1988; Clarke, Bellmann, Meuli, Assal, & Steck, 2000; Bunzeck, Wuestenberg, Lutz, Heinze, & Jancke, 2005).

Finally, these results serve as essential evidence to suggest that aphantasia is a valid experience, defined by the inability to form voluntary visual images with a selective impairment in object imagery. Previous work has shown relatively intact performance by aphantasics on imagery and visual working memory tasks (Jacobs et al., 2018), and some researchers have proposed aphantasia may be more psychogenic than a real impairment (de Vito & Bortolomeo, 2016). However, in the current study, we observe evidence for a selective impairment in object imagery for aphantasics in comparison to controls. Importantly, if such an impairment were caused by intentional efforts to demonstrate an impairment, we would expect decreased performance in spatial accuracy, decreased performance in the perceptual drawing task, or low ratings in all questions of the OSIQ rather than solely the object imagery component. However, in all of these cases, aphantasics performed identically with controls. In fact, aphantasics even showed higher memory precision than controls on some measures, including significantly fewer memory errors and fewer editing in their drawings. Further, the correlations between the VVIQ, OSIQ, and drawn object information lend validity to the self-reported questionnaires in capturing true behavioral deficits. This being said, while we observed a deficit in object memory for aphantasics, it was not a complete elimination of object memory abilities. Aphantasics were still able to draw five objects per image from memory. While this moderate performance could be due to some preserved ability at object memory, this performance could also reflect the use of verbal lists of objects combined with intact, accurate spatial memory to reconstruct a scene. Future work will need to directly compare visual and verbal strategies, and push the limits to see what occurs when there is more visual detail than can be supported by verbal strategies.

In conclusion, leveraging the wide reach of the internet, we have conducted an in-depth and large scale study of the nature of aphantasics’ mental representations for visual images. Aphantasics have a unique mental experience that can provide essential insights into the nature of imagery, memory, and perception. Their drawings reveal a complex, nuanced story that show impaired object memory, with a combination of semantic and spatial strategies used to reconstruct scenes from memory. Collectively, these results suggest a dissocation in object and spatial information in visual memory.

## Acknowledgements

This research was supported (in part) by the Intramural Research Program of the National Institute of Mental Health (ZIA-MH-002909).

